# Sequencing using a two-steps strategy reveals high genetic diversity in the S gene of SARS-CoV-2 after a high transmission period in Tunis, Tunisia

**DOI:** 10.1101/2021.06.18.449083

**Authors:** Wasfi Fares, Kais Ghedira, Mariem Gdoura, Anissa Chouikha, Sondes Haddad-Boubaker, Marwa Khedhiri, Kaouthar Ayouni, Asma Lamari, Henda Touzi, Walid Hammemi, Zina Medeb, Amel Sadraoui, Nahed Hogga, Nissaf ben Alaya, Henda Triki

## Abstract

Recent efforts have reported numerous variants that influence SARS-CoV-2 viral characteristics including pathogenicity, transmission rate and ability of detection by molecular tests. Whole genome sequencing based on NGS technologies is the method of choice to identify all viral variants; however, the resources needed to use these techniques for a representative number of specimens remain limited in many low and middle income countries. To decrease sequencing cost, we developed a couple of primers allowing to generate partial sequences in the viral S gene allowing rapid detection of numerous variants of concern (VOCs) and variants of interest (VOIs); whole genome sequencing is then performed on a selection of viruses based on partial sequencing results. Two hundred and one nasopharyngeal specimens collected during the decreasing phase of a high transmission COVID-19 wave in Tunisia were analyzed. The results reveal high genetic variability within the sequenced fragment and allowed the detection of first introduction in the country of already known VOCs and VOIs as well as others variants that have interesting genomic mutations and need to be kept under surveillance.

**Importance:** The method of choice for SARS-CoV-2 variants detection is whole genome sequencing using NGS technologies. Resources for this technology remain limited in many low and middle income countries where it is not possible to perform whole genome sequencing for representative number of SARS-CoV-2 positive cases. In the present work, we developed a novel strategy based on a first partial sanger screening in the S gene including key mutations of the already known VOCs and VOIs for rapid identification of these VOCs and VOIs and helps to better select specimens that need to be sequenced by NGS technologies. The second step consisting in whole genome sequencing allowed to have a holistic view of all variants within the selected viral strains and confirmed the initial classification of the strains based on partial S gene sequencing.

## Introduction

Severe acute respiratory syndrome coronavirus 2 (SARS-CoV-2), which is the causative agent of human coronavirus disease 2019 (COVID-19), was identified in Wuhan-China in December 2019 (1, 2). The outbreak of the coronavirus disease (COVID-19) rapidly spread worldwide; it was officially declared as pandemic by the World Health Organization (WHO) on March 11, 2020 (3) and now represents a tremendous threat globally.

SARS-CoV-2 is a single-stranded positive RNA virus, a member of the Beta coronavirus genus that also contains SARS-CoV and MERS-CoV. The first sequence of the virus was published in January 2020 (4). The structural genome region, located in the 3’ part of the genome, encodes four structural proteins: spike (S), envelope (E), membrane (M) and nucleocapsid (N) (5). The S protein forms a trimer on the surface of the virion, it mediates virus attachment to the ACE-2 receptor and its entry to the host cells (6). The S Protein is composed of two sub-units, S1 containing the receptor-binding domain (RBD) and S2 that mediates membrane fusion (7). The S protein determines SARS-CoV-2 infectivity and transmissibility and is also the major antigen inducing protective immune response (8). Since the beginning of the COVID-19 pandemic, the S protein has been undergoing several mutations and it is highly important to follow the emergence of these variants and their biological, epidemiological and clinical significance. Early in the pandemic, variants of SARS-CoV-2 containing a D to G substitution in the 614 amino-acid residue of the S protein (D614G) were reported. This substitution increased receptor binding avidity and D614G mutants became dominant in many geographic regions (9–11). In December 2020, the United Kingdom reported a variant of concern (VOC), referred as B.1.1.7, with enhanced transmissibility within the population (12, 13). This variant became predominant in the UK and spread to more than 100 countries in the world. In January 2021, two other VOCs, referred as B.1.351 and B1.1.28, also with high transmissibility, were reported in South Africa and Brazil, respectively (14–16). Later, many other variants, classified as Variants Under Investigation (VUIs) were reported throughout the world. In addition to the increased transmissibility, it is suggested that some mutations in these variants may affect the performance of some diagnostic real-time PCR tests and reduce susceptibility to vaccine-induced neutralizing antibodies (9, 10, 17–22). Global tracking of these newly identified VOCs and VUIs as well as any other evolving SARS-CoV-2 variant, by genomic surveillance and rapid sharing of viral genomic sequences, is highly recommended in order to limit their spread and control the pandemic.

Nowadays, several classifications of SARS-Co V-2 strains in lineages or clades were proposed. Indeed, two different lineages, A and B, were proposed by the Phylogenetic Assignment of Named Global Outbreak (PANGO) lineage nomenclature, while a classification in 11 different clades (19-A, 19-B, 20-A to 20-I) was proposed by the Nextstrain resources and another classification in 9 clades (S, L, O, V, G, GH, GR, GRY and GV) was proposed by GISAID.

In Tunisia, the first case of SARS-CoV-2 infection was reported on March 03, 2020 (23). The country experienced a first wave of the COVID disease and, through setting up drastic nation-wide multi-sectoral measures to avoid international introduction of the virus and its spread within the population, COVID-19 incidence decreased in May-June 2020 to reach zero cases per day from the 4th to the 11th of June 2020. The national strategy included early detection of imported cases, quarantining of new confirmed cases as well as suspected cases and strict travel restrictions. After the sharp decrease of the disease incidence; a relaxation in the application of these measures by the general population, combined with decreased restrictions in international transportation, led to the re-introduction of the virus again and the establishment of a local transmission. In late July, COVID-19 incidence started to increase again and the country experienced a second wave with highest incidence in January 2021, associated with a high local transmission within the population. Starting from February 2021, the disease incidence together with mortality rates decreased again.

The present work reports the genomic features of SARS-CoV-2 sequences detected in Tunisia during the late phase of the second wave of the pandemy and reveals the co-circulation of several variants, some of which are already known as VOCs, others have interesting genomic mutations and need to be kept under surveillance.

## Material and Methods

### Nasopharyngeal samples

A total of 201 SARS-CoV-2 positive nasopharyngeal samples, collected from individuals living in the four districts of Tunis capital, were included in this study. Sample collection was performed from January to March 2021, during the decreasing phase of the second wave of COVID-19 outbreak in Tunisia (**Figure 1**). The study population includes symptomatic patients presenting with mild COVID clinical forms or with severe forms as well as asymptomatic individuals sampled after a contact with confirmed cases. The study population included 91 males and 110 females, their age ranged from 5 to 98 years. The samples were collected by the health teams from the Ministry of Health, at home for asymptomatic individuals and those with non-severe clinical symptoms, or at the health facility level for hospitalized patients. Samples were transported, refrigerated and within 24 hours, to the Pasteur Institute of Tunis where they were immediately processed for SARS-CoV-2 detection by specific real time reverse transcription polymerase chain reaction (RT-PCR) according to WHO approved protocols (24, 25).

**Figure 1.**
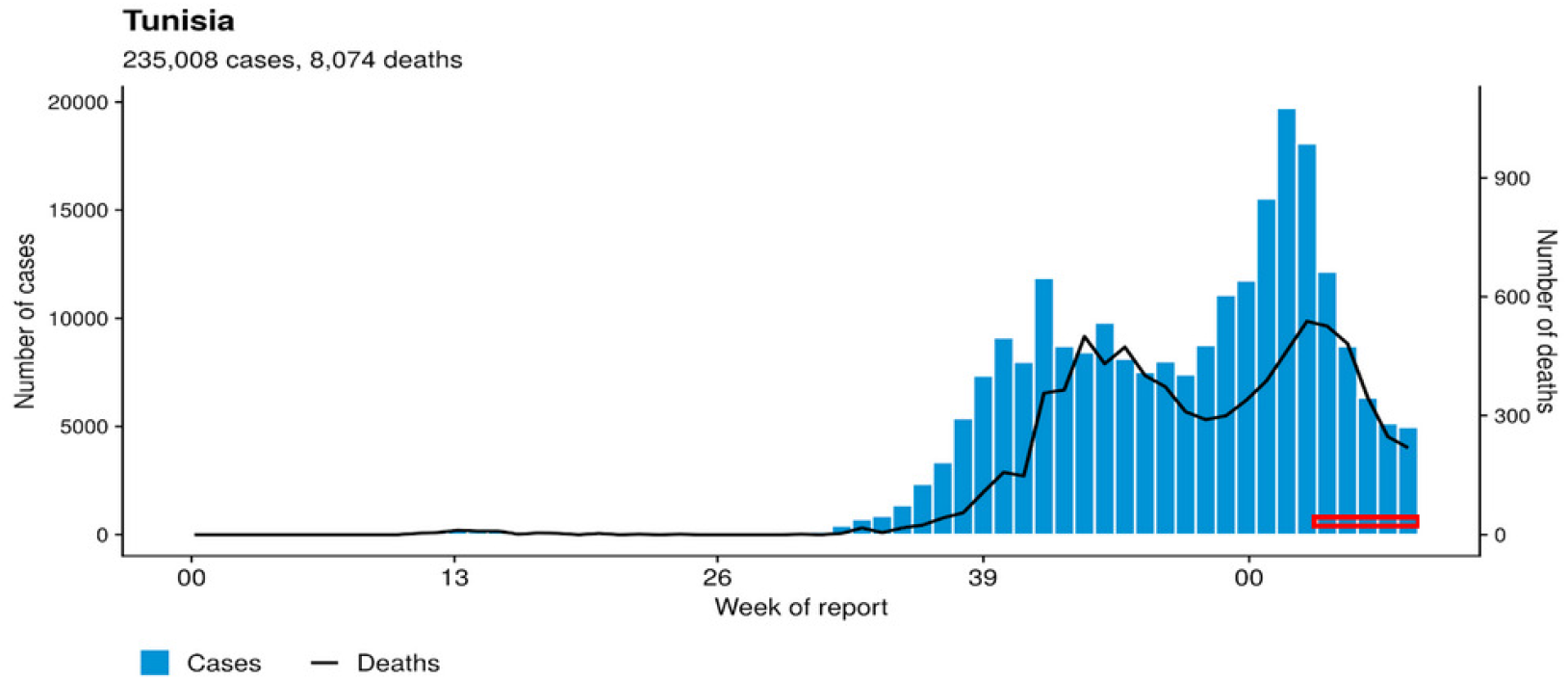
Samples collection period investigated in the present study. The graph displays the number of cases and the number of deaths in Tunisia since the declaration of the pandemy in March 2020. The Abscisse axe represents the number of weeks from March 2020 till May 2021. Weeks highlighted in red color represents the samples collection period investigated in the present study.

### Ethical statement

This work was performed in the frame of COVID-19 diagnostic effort, and all samples used for analysis were anonymized. This study was approved by the Bio-Medical Ethics Committee of the Pasteur Institute of Tunis, Tunisia réf. 2020/14/I/LR16IPT/V1

### Primer design

Primers were designed using PrimerDesign-M online software, available through https://www.hiv.lanl.gov/content/sequence/PRIMER_DESIGN/primer_design.html (26, 27), based on an alignment of 13451 SARS-CoV-2 complete genome sequences. Several points were considered such as melting temperatures, G+C percentage, entropy, complexity and nucleotide composition, in order to perfectly align with the SARS-CoV-2 sequence. The selected primers sequences were as follows: IPT_FW: (22964-22987) 5’-ATTTCAACTGAAATCTATCAGGCC-3’ and IPT_REV: (23666-23647) 5’-CTGCACCAAGTGACATAGTG-3’. Indicated positions correspond to the sequence of Wuhan reference strain (accession number: NC045512). The designed primers allow the amplification of a 703-nucleotide-long region in the S gene holding key mutations, that includes the E484K, N501Y, A570D, D614G and P681H, recently identified as specific of the main VOCs and VUIs of SARS-CoV-2.

### PCR amplification and sequencing in the S gene

A volume of 140μl of nasopharyngeal samples was used for viral RNA extraction with viral RNA Mini Kit (Qiagen, Hilden, Germany) to give a final elution volume of 60μl of total RNA. The presence of SARS-CoV-2 RNA was determined by conventional reverse transcription PCR using the SuperScript^®^III One-Step RT-PCR System with Platinum^®^ Taq DNA Polymerase kit (Invitrogen) in a 25μl reaction volume containing 12.5μl of 2X buffer, 0.5μl Rnasin (Promega), 1 μl of each reverse and forward primers (10μM), 1μl Enzyme mix and 5μl of total extracted RNA. Optimized cycling conditions was performed as follows: Reverse transcription with initial incubation at 50°C for 30min and 94°C for 2min followed by 35 cycles, repeating denaturation at 94°C for 15sec, annealing at 54°C for 45sec and elongation at 72°C for 30sec, and final elongation at 72°C for 10min. Amplification products are first visualized by electrophoresis in agarose gels and then purified by the ExoSAP-IT method using the Exonuclease-I and the Shrimp Alkaline Phosphatase (Invitrogen). The purified amplicons were sequenced using the Big Dye Terminators v3.1 kit (Applied Biosystems) and the forward and reverse PCR primers. The resulting consensus sequences were deduced by aligning the forward and the reverse sequence of each isolate, excluding primer binding regions and are 618 nucleotides-long (positions 22988 to 23605 according to the Wuhan reference strain NC045512). They were submitted to the NCBI database under accession number MZ150010 - MZ150210.

### Whole genome sequencing

The QIAseq SARS-CoV-2 Primer Panel paired with the QIAseq FX DNA Library construction kits (Qiagen GmbH, Germany) were used for enriching and sequencing the entire SARS-CoV-2 viral genome. Extracted RNA from nasopharyngeal swabs was first depleted of ribosomal RNA using RiboZero rRNA removal Kit (Illumina, USA). The residual RNA was then converted to double stranded cDNA using random priming. Following cDNA synthesis, QIAseq SARS-CoV-2 Primer Panel kit was used including high fidelity multiplex PCR reaction yielding 400bp amplicons covering the full viral genome. The multiplexed amplicon pools were then converted to sequencing libraries by enzymatic fragmentation with 250bp fragment size, end repair and ligation to adapters with the QIAseq FX DNA Library construction kits. Thereafter, the constructed DNA library was purified and adapter-dimers were removed with Agencourt AMPure XP beads. The libraries were sequenced using Nextseq (Illumina Inc, USA) to generate 2×150 bp paired-end sequencing reads.

Sequences’ raw data have been processed using fastqc version 0.11.9 for quality control (**https://www.bioinformatics.babraham.ac.uk/projects/fastqc/**). Low quality reads and adapters have been filtered using trimmomatic version 0.39 (28) with a Phred quality score of 30 as threshold. Genome consensus sequences were assembled by mapping on the SARS-CoV-2 reference genome of GenBank accession number NC045512 (Wuhan-Hu-1 isolate) using Spades assembler version 3.15.0 (29), with thresholds of 80% for nucleotide sequence coverage and 90% for nucleotide similarity. The obtained SARS-CoV-2 new sequences were submitted to the GISAID database (https://www.gisaid.org) (30, 31) with the following accession numbers: EPI_ISL_2035560, EPI_ISL_2035563, EPI_ISL_2035720, EPI_ISL_2035734, EPI_ISL_2035752, EPI_ISL_2035753, EPI_ISL_2035940 to EPI_ISL_2035949, EPI_ISL_2035988 and EPI_ISL_2036077.

### Phylogenetic analysis

The obtained partial S gene sequences and selective whole genome sequences were aligned together with representative SARS-CoV-2 reference sequences of the nine recognized GISAID clades publically available in the GISAID database using MUSCLE multiple sequence alignment algorithms (32) implemented in MEGAX (33). Phylogenetic analyses were performed on nucleotide sequences using the maximum likelihood method with the Tamura 3-parameter model then on amino acid sequences, obtained from the aligned sequences, using the maximum likelihood method and the Jones Taylor Thornton model. The tree topologies were supported by 1000 bootstrap replicates.

Mutation profiles in the ORF1a, ORF1b, S, ORF3a, E, M, ORF6, ORF7a, ORF8, N, and ORF10 genomic regions of SARS-CoV-2 were assessed, by comparing the nucleotide and deduced amino acid sequences of the Tunisian strains with those of the Wuhan reference strain, using the sequence alignment performed by MUSCLE multiple sequence alignment algorithms (32) implemented in MEGAX (33).

## Results

Phylogenetic tree in **Figure 2** was performed based on the alignment of the 618-nucleotides fragment in the S gene of the 201 studied Tunisian SARS-CoV-2 strains, together with the 9 selected references SARS-CoV-2 sequences according to the GISAD nomenclature. The tree topology shows that the Tunisian sequences are divided into 3 different clusters. Cluster1, represented in purple color, includes the highest number of sequences (174 out of 201,86.5% of Tunisian strains) that clustered with the 4 reference sequences of the GISAID Clades G, GH, GR and GV. The phylogenetic distribution within this cluster shows several phylogenetic sub-branches reflecting a large genetic variability. Cluster2 indicated in blue color comprises 15 identical sequences that clustered with the GISAID reference sequence from Clade GRY. Cluster3, indicated in red color, contains 12 sequences that clustered with the GISAID reference sequence from Clade S.

**Figure 2.**
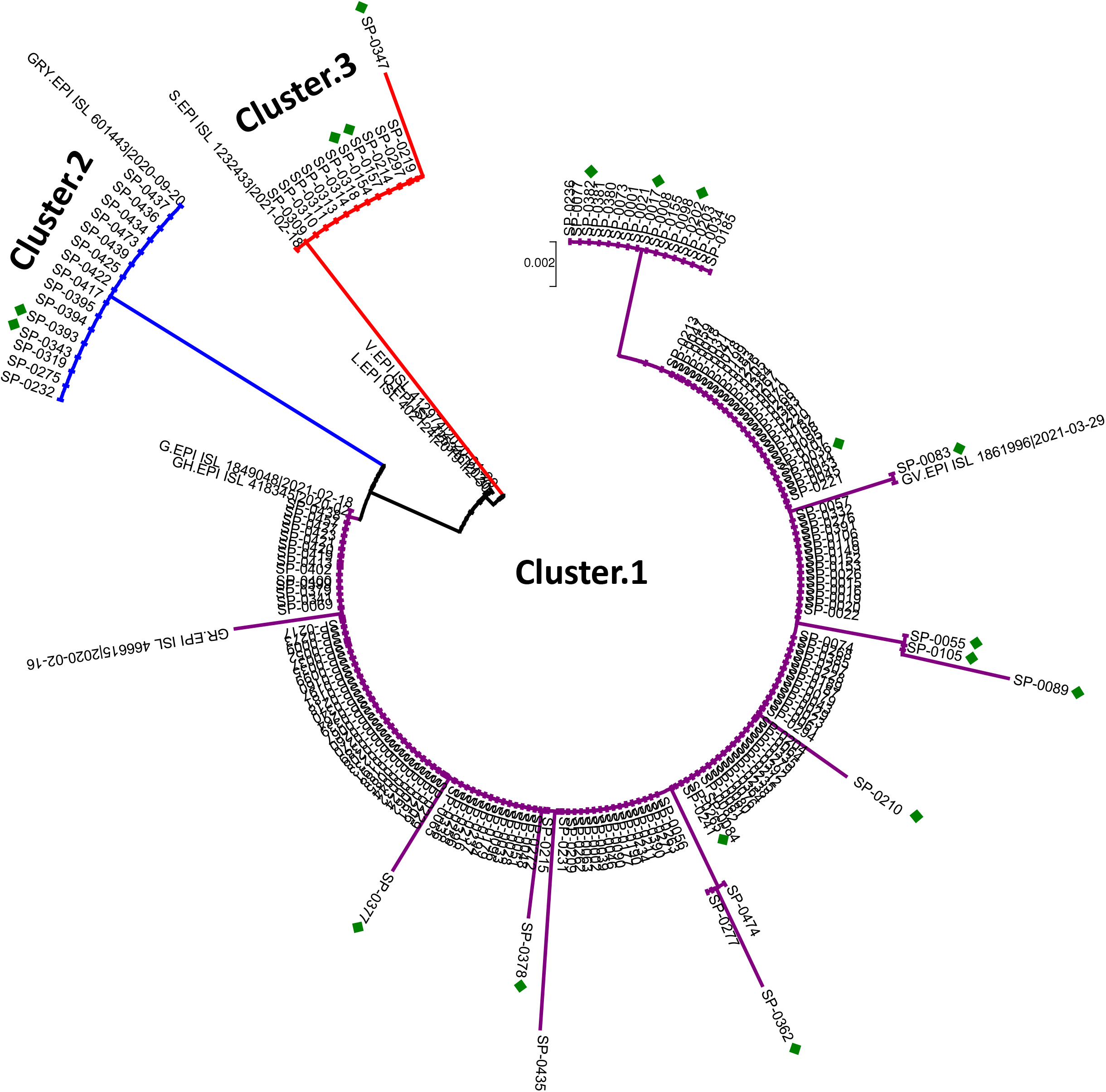
Phylogenetic tree of 201 SARS-CoV-2 sequences based on partial S gene nucleotide sequencing. The phylogenetic tree includes 201 Tunisian sequences compared to 9 representative reference sequences of SARS-Cov-2 Clades. The tree was performed using the neighbor joining method and the Tamura 3-parameter (T92) model. Topology was supported by 1000 bootstrap replicates. The sequences reported in this study are shown in bold, and indicated by the laboratory code. The sequences downloaded from GISAID are indicated by their accession number followed. Cluster 1 in purple color denotes sequences presenting the D614G substitution and the lack of the amino acid substitution N501Y. Cluster 2 in blue color includes sequences having the N501Y, A570D, D614G and P681H substitutions. Cluster 3 in red color groups sequences with the N501Y, A653V and H655Y substitutions and the lack of the amino acid substitution D614G.

Eighteen representative samples from these clusters, indicated by a green square in Figure 2, were selected for whole genome sequencing: 13 from Cluster1, 2 from Cluster2 and 3 from Cluster3. The phylogenetic tree of the obtained 18 whole genome sequences, together with the 9 GISAID references SARS-CoV-2 sequences, is shown in **Figure 3**. The figure also shows the classification of the Tunisian sequences according to the PANGO and the Nextstrain classifications. The phylogenetic distribution of the sequences based on whole genome sequences (Figure 3) is similar to the one obtained in Figure 2, based on the partial S gene genomic data.

**Figure 3.**
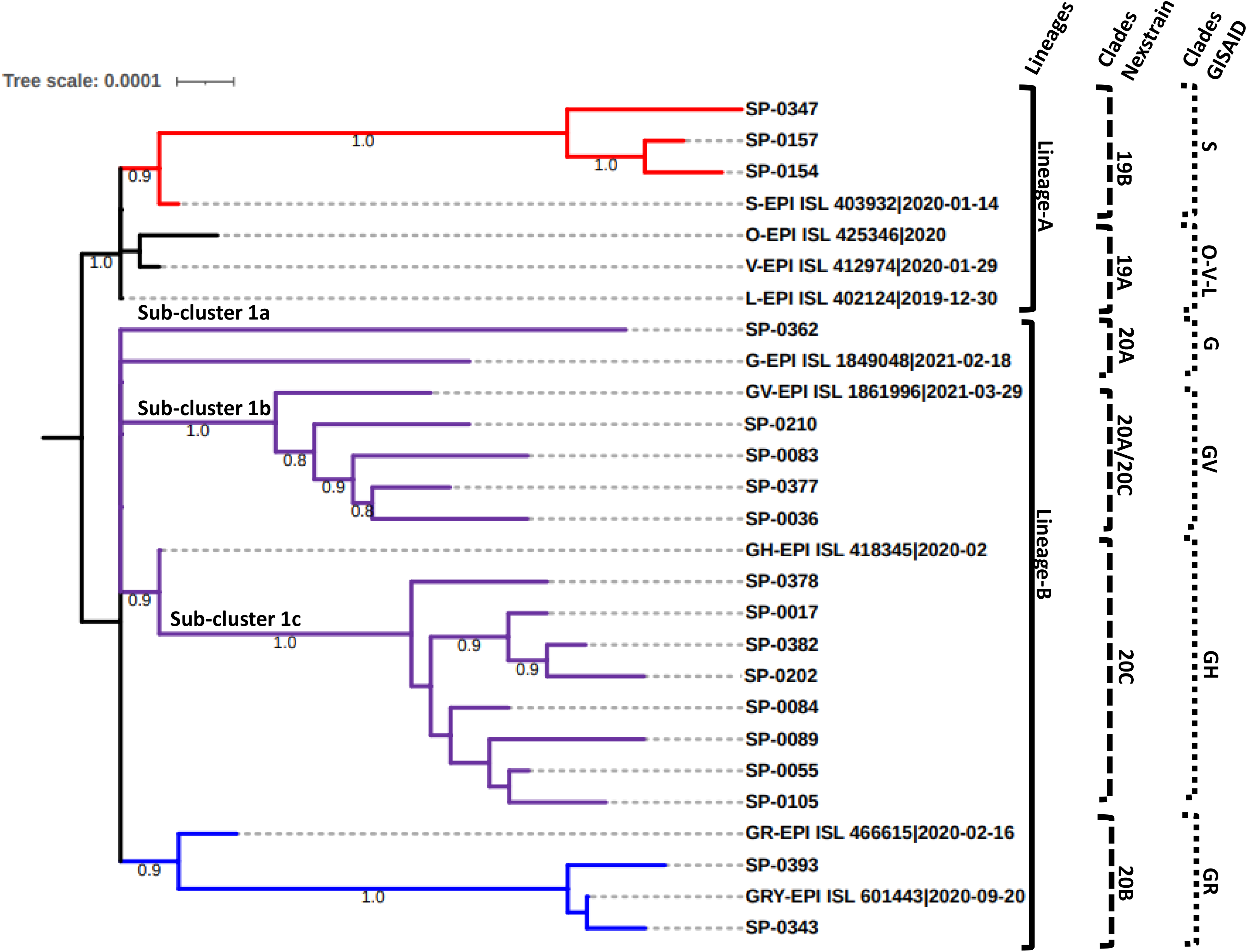
Phylogenetic tree of 18 SARS-CoV-2 whole genome sequences circulating in Tunisia compared to 9 reference strain genomes. The phylogenetic tree includes 18 Tunisian sequences compared to 9 representative reference sequences of SARS-Cov-2 Clades. The tree was performed using the neighbor joining method and the Tamura 3-parameter (T92) model. Topology was supported by 1000 bootstrap replicates. The sequences reported in this study are shown in bold, and indicated by the laboratory code. The sequences downloaded from GISAID database are indicated by their accession number. Cluster 1 in purple color, Clade 2 in blue color and Clade 3 in red.

The 13 sequences from Cluster1 highlighted in purple color in Figure 2 grouped together within PANGO B lineage in Figure 3. The phylogenetic distribution of these sequences clearly shows the presence of 3 sub-clusters called sub-cluster 1a, 1b and 1c classified as clade G/20A, GV/20A-C and GH/20C respectively according to the GISAID/Nextstrain nomenclatures. Sub-cluster 1a is represented by only one sequence (SP-0362), while Sub-cluster 1b and Sub-cluster 1c are represented by 4 (SP-0202, SP-0083, SP-0377 and SP-0036) and 8 (SP-0378, SP-0017, SP-0382, SP-0210, SP-0084, SP-0089, SP-0055 and SP-0105) sequences respectively.

Two sequences from Cluster2 in Figure 2 were also found to cluster together with the reference sequence of the GR GISAID Clade based on whole genome sequencing comparison; the sequences also belong to the PANGO B lineage and to the 20B Clade of the Nextstrain nomenclature.

Unlike the sequences from Cluster1 and Cluster2, the three whole genome sequences from Cluster3 belong to the PANGO A lineage. They grouped together with the reference sequence of the S GISAID Clade, similarly to the results obtained based on the partial S sequences.

The amino acid sequences related to the 201 partial S sequences and the 18 whole genome sequences were deduced from the obtained nucleotide sequences and compared to the Wuhan reference protein sequences.

Table 1 shows the amino acid substitution profile in the sequenced fragment of the S gene of the 201 samples investigated in the present study. Fourteen different mutation profiles were found. Most of the sequences (147/174) had zero non synonymous mutation as compared to the Wuhan reference, excepting the D614G which was found in all the sequences from cluster 1 and Cluster 2. The remaining 27 sequences from Cluster1 had 1 to 2 additional substitutions within the sequenced fragment (**Table 1**). The 15 sequences from Cluster 2 shared an identical mutational profile with the amino acid substitutions N501Y, A570D, D614G and P681H which are known to be characteristic of the VOC B.1.1.7 initially detected in the UK. The 12 sequences from Cluster did not have the D614G substitution but three mutations that suggest the VUI A.27 (N501Y, A653V, Q655H); one sequence (SP-0347 which was in a separate branch within the phylogenetic tree shown in Figure 2), had an additional substitution (Q677H).

**Table. 1.**
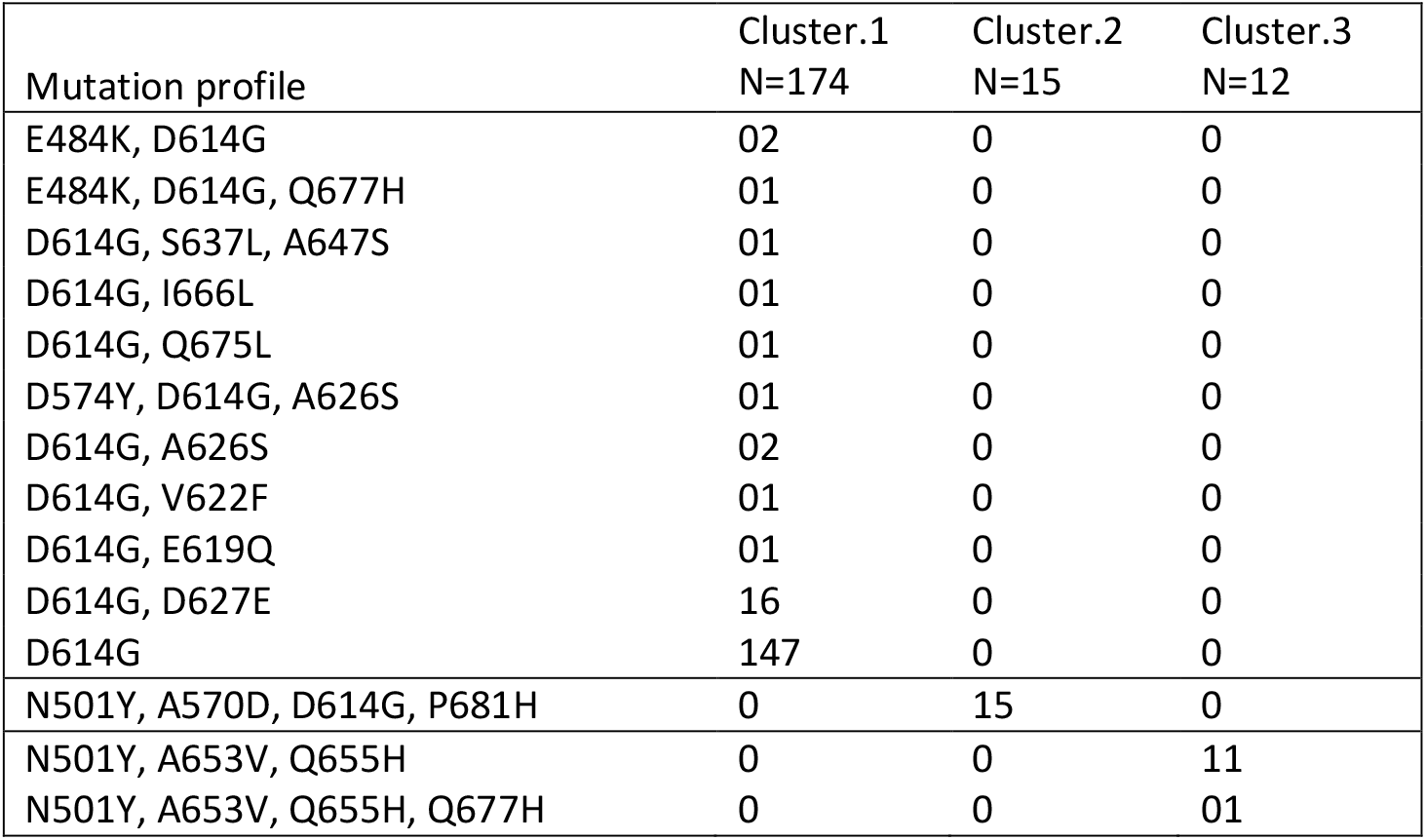
Amino acid substitution profile in the sequenced fragment of the S gene

Table 2 shows the amino acid substitution profile along the whole genome of the 18 selected Tunisian SARS-CoV-2 and representative from the different clusters found based on S partial sequences. The two sequences from Cluster 2 had identical mutational profile in the S gene and a total of 23 and 24 amino acid substitution along the whole genome; these results confirm the belonging of the two sequences to the B1.1.7 lineage (VOC). The three sequences from Cluster 3 shared 15 identical amino acid substitutions along the whole genome and the results confirm the belonging of the three sequences to the A.27 lineage, identified as variant of interest (VOI) initially detected in France. Among Cluster 1, one sequence (SP062 - Sub-Cluster 1a) had a mutational profile that corresponds to the identified variant of interest (VOI) B.1.525 initially detected in Nigeria and in the UK. The sequences from Sub-Cluster 1c shared several identical mutations in the non structural regions of the genome and belonged to the B.1.160 lineage that is not presently identified as VOC or VOI. The same is for the sequences from Sub-Cluster 1b that had more genetic diversity and belonged to the B.1.177 lineage.

**Table 2:**
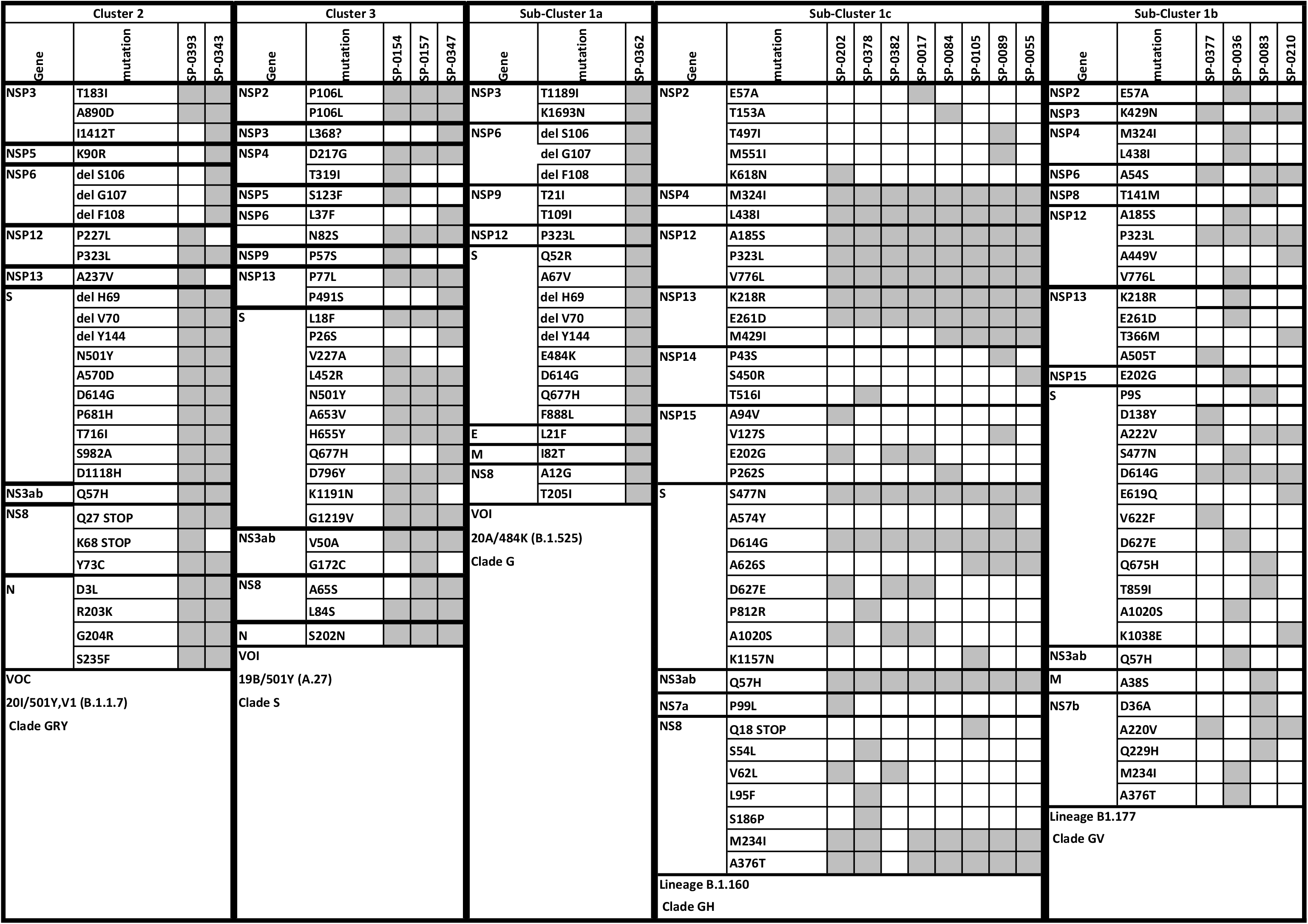
Mutation profile of the 18 obtained Sars-Cov-2 Tunisian strains

## Discussion

Since the beginning of the COVID-19 pandemic, several SARS-CoV-2 variants emerged, some of them totally changed the infection epidemiology. First, a variant with the D614G mutation emerged and became dominant globally (34). In our series, this mutation is found in 186 out of the 201 isolates (92%). Other variants emerged subsequently and it is now hypercritical to track the already known of them labelled as VOCs or VOIs and also to monitor the emergence of new variants. The method of choice is whole genome sequencing using NGS high throughput technologies which improved considerably during the last years with a cost that declines continuously. However, despite this progress, NGS is still expensive and resources for this technology remain limited in many low and middle income countries where it is not possible to perform whole genome sequencing for representative number of SARS-CoV-2 positive cases. Thus, the use of other technologies to identify isolates that are to be sequenced in priority is highly needed. Several real-time PCR tests that target the already known VOCs, especially the B.1.1.7 (United Kingdom), B.1.351 (South Africa) and B1.1.28 (Brazil) are now commercially available. They can be very useful to rapidly identify the introduction of these VOCs to a country/region and to monitor their transmission. However, these kits cannot detect other variants of interest that already emerged, or that may emerge any time. Furthermore, other variants can be characterized by the failure to detect the S gene in these tests, known as S gene target failure (SGTF) (35).

In the present work, we developed a couple of primers allowing to generate a 618-nucleotide-long sequence in the viral S gene that includes key mutations of the already known VOCs and VOIs. Sequencing of this fragment by the traditional Sanger technology allows rapid identification of these VOCs and VOIs and helps to better select specimens that need to be sequenced by NGS technologies. Using this approach, it is possible to detect 14 amino acid substitutions that have been identified in several VOCs and VOIs (G482V, E484K, N501Y, A570D, D574Y, D614G, E619Q, A626S, D627E, A653V, H655Y, Q675H, Q677H and P681H) and to get a rapid orientation towards an already known or a new variant. In our series and using these primers, we were able to detect the first introduction of the B.1.1.7 (VOC) and two other VOIs (A.27 and B.1.525) and to select other viruses for WGS based on the results obtained in the S partial genomic region. The second step consisting in whole genome sequencing allowed to have a holistic view of all variants within the selected viral strains and confirmed the initial classification of the strains based on partial S gene sequencing.

The specimens included in the present work were collected in the decreasing phase of a COVID-19 wave that occurred in Tunisia starting from September 2020 up to January 2021. This period was characterized by a high transmission within the population and this explains the high genetic diversity that we found in the obtained sequences. Several lineages were identified and more than 100 different amino acid changes, in comparison to the standard Wuhan strain, were identified all through the viral genome.

During the study period, the first isolates of the VOC B.1.1.7, initially identified in the UK, were detected. The sequenced isolates had the H69del, V70del, Y144del, N501Y, A570D, D614G P681H, T716I, S982A, D1118H common amino acid substitutions with the 20I/501Y.V1 (UK variant). Thus, it is highly expected that the genetic features described herein will rapidly change to a lower genetic variability and a predominance of the B.1.1.7 UK lineage. Indeed, this is what happened in most countries of the world where the B.1.1.7 UK lineage was introduced causing devastating waves of COVID-19 (36, 37). With its higher transmissibility within the human population, it becomes rapidly predominant once introduced and this is also what is expected to happen in Tunisia.

Furthermore, we were able to detect viruses belonging to the A.27 lineage, initially detected in Danemark and now classified as VOI. This lineage was detected in around 26 different countries in the world, from Europe, Africa as well as USA and Australia.

Whole genome sequencing of three isolates in this series revealed the presence of amino acid substitutions characteristic of this lineage, including L18F, L452R, N501Y, A653V, H655Y, D796Y and G1219V and the absence of the D614G substitution in the Spike protein. One strain (SP-0347) presented two additional substitutions: P26S that is found in the P1 20J/501Y.V3 (Brazilian variant) and Q677H found in the *Henri Mondor variant* detected in different regions of France (38).

We have also detected one sequence (SP062 - Sub-Cluster 1a) with a mutational profile corresponding to the B.1.525, initially detected in Nigeria and in the UK. This variant was detected in 48 different countries in the world at writing time and is presently classified as VOI.

The rest of sequences from Sub-Clusters 1b and 1c belonged to the B.1.160 and B.1.177 lineages that are not presently identified as VOCs or VOIs. These sequences exhibit quite high genetic variability which is expected after the high active transmission period that the country experienced in late 2020 and January 2021. Among all these variants, some may then disappear and other may persist or even dominate if they have a selective advantage in terms of virulence or transmissibility.

## Conclusion

In conclusion, this study gives an overview of the SARS-CoV-2 strains circulating in Tunisia after a high transmission wave of COVID-19. Partial S gene sequencing followed by whole genome sequencing of a selection of specimen was used to identify the different circulating variants. This strategy may be of interest for several countries; it helps to establish a genomic surveillance that is now highly needed in all regions of the world, with a good cost/effectiveness ratio.

## ACKNOWLEDGMENTS

This work was co-founded by the Tunisian Ministry of High education and Research and the European Union’s Horizon 2020 research and innovation program under grant agreement No 883441, project STAMINA (Demonstration of intelligent decision support for pandemic crisis prediction and management within and across European borders). The authors thank the field staff from the Ministry of Health for their efforts in specimen collection and transportation to the laboratory.

